# Heartbeat and Somatosensory Perception

**DOI:** 10.1101/2020.12.29.424693

**Authors:** Esra Al, Fivos Iliopoulos, Vadim V. Nikulin, Arno Villringer

**Affiliations:** Department of Neurology, Max Planck Institute for Human Cognitive and Brain Sciences, 04103 Leipzig, Germany; MindBrainBody Institute, Berlin School of Mind and Brain, Humboldt-Universität zu Berlin, 10099 Berlin, Germany; Center for Stroke Research Berlin (CSB), Charité – Universitätsmedizin Berlin, 10117 Berlin, Germany; Institute of Cognitive Neuroscience, National Research University Higher School of Economics, 101000 Moscow, Russia

**Keywords:** Somatosensory processing, perceptual awareness, electrophysiology, interoception

## Abstract

Our perception of the external world is influenced by internal bodily signals. For example, we recently showed that timing of stimulation along the cardiac cycle and spontaneous fluctuations of heartbeat-evoked potential (HEP) amplitudes influence somatosensory perception and the associated neural processing (Al et al., 2020). While cardiac phase affected detection sensitivity and late components of the somatosensory-evoked potentials (SEPs), HEP amplitudes affected detection criterion and both early and late SEP components. In a new EEG study, we investigate whether these results are replicable in a modified paradigm, which includes two succeeding temporal intervals. Only in one of these intervals, subjects received a weak electrical finger stimulation and then performed a yes/no and two-interval forced-choice detection task. Our results confirm the previously reported cardiac cycle and prestimulus HEP effects on somatosensory perception and evoked potentials. In addition, we obtain two new findings: A source analysis in these two studies shows that the increased likelihood of conscious perception goes along with HEP fluctuations in parietal and posterior cingulate regions, known to play important roles in interoceptive processes. Furthermore, HEP amplitudes are shown to decrease when subjects engage in the somatosensory task compared to their resting state condition. Our findings are consistent with the view that HEP amplitudes are a marker of interoceptive (versus exteroceptive) attention and provide a neural underpinning for this view.

## 1. Introduction

Interoceptive signals from the body can influence our perception, cognition, and emotions (Critchley & Garfinkel, 2018; Garfinkel et al., 2014; Sandman, McCanne, Kaiser, & Diamond, 1977; Saxon, 1970). For instance, different phases of the cardiac cycle (e.g., systole vs. diastole) have been reported to affect visual and auditory perception (Sandman et al., 1977; Saxon, 1970). In the somatosensory domain, we recently demonstrated that somatosensory detection decreases during the systole compared to the diastole phase of the cardiac cycle in two independent studies (Al et al., 2020; Motyka et al., 2019 but also see Edwards, Ring, Mcintyre, Winer, & Martin, 2009). Furthermore, our latest study (Al et al., 2020) showed that this change in detection went along with changes in perceptual sensitivity for discriminating the presence of stimuli from its absence, rather than in detection bias, i.e., criterion, according to signal detection theory (SDT). Similar to the changes in perception, late components (P300) of somatosensory-evoked potentials (SEPs) were found to be lower during systole compared to diastole (Al et al., 2020). We explained these findings using an interoceptive predictive framework, which suggests that rhythmic bodily signals such as heartbeat-related events are predicted, therefore suppressed from entering conscious perception and that the same suppressive mechanisms also inhibit perception of coincident weak external stimuli.

Our previous study also addressed the influence of neural responses to heartbeats, known as heartbeat-evoked potentials (HEP), on somatosensory perception. Higher amplitudes of prestimulus HEP were found to be followed by lower detection rates and decreased amplitudes of early (P50) and late SEP components (N140, P300) (Al et al., 2020). The modulation of detection rates was associated with a change in detection bias but not sensitivity. Given that HEP amplitudes are higher during interoceptive compared to exteroceptive attention (García-Cordero et al., 2017; Petzschner et al., 2019; Villena-González et al., 2017), we interpreted the fluctuations of HEP amplitudes as a reflection of attentional shifts between internal bodily states and external somatosensory stimuli.

The main objective of the current study was to replicate the previous findings (Al et al., 2020) in a different experimental setting which included two subsequent time intervals. Based on our previous findings, we hypothesized (i) a lower detection sensitivity and weaker late SEP amplitudes during systole as compared to diastole and (ii) negative relationships between HEP amplitudes and succeeding detection as well as the amplitudes of early and late SEP components. Beyond the replication of our previous study, we attempted to identify (iii) the neural sources of HEP differences preceding hits and misses and we investigated (iv) whether HEP amplitudes differ during a somatosensory detection task compared to a resting state condition.

## 2. Materials and Methods

### 2.1 Participants

Forty healthy volunteers participated in the study. Four subjects were excluded from the analysis due to technical problems during the ECG data acquisition. Overall, 252 experimental blocks with 30240 trials in 36 subjects (18 females, age: 25.3 ± 4.0 y [mean ± SD], range: 20 to 36 y) were included in the analyses.

### 2.2 Ethics statement

The study was approved by the ethics commission at the medical faculty of the University of Leipzig (no. 462-15-01062015). All participants gave informed consent and were compensated for their participation.

### 2.3 Somatosensory Stimulation and Task Design

Somatosensory stimuli were presented using electrical stimulation of the finger nerve. A pair of steel wire ring electrodes was attached to the middle (anode) and the proximal (cathode) phalanx of the left index finger. Participants sat comfortably in a quiet room, with palms facing downward. Stimuli were applied using a DS-5 constant-current stimulator (Digitimer) using single square-wave pulses with a duration of 200 μs.

At the beginning of the experiment, a near-threshold stimulus intensity that could be detectable only in 50% of the trials in the yes/no detection task was determined by using a two-step procedure as described in our previous study (Al et al., 2020).

During the experiment, participants performed two detection tasks: a yes/no detection task and a two-interval forced-choice detection task (2IFC) in every trial. Each trial was separated into two temporal intervals and each interval had a duration of 1400 ms (**Fig. 1**). In stimulation trials, participants either received an electrical stimulus during the middle of the first or the second temporal interval. After the visual representation of the second interval disappeared from the screen, participants first answered the question, ‘did you feel it?’, as quickly as possible by reporting ‘yes’ or ‘no’ with their right middle finger using button presses. Subjects were then asked to respond to the question, ‘in which temporal interval did the stimulus occur?’, by choosing either ‘1’ with their right index finger or ‘2’ with their right ring finger. They were instructed ‘to guess’ the interval even if they reported not detecting the stimulus presence. Participants were told that one of the two temporal windows contained a stimulus in every trial; however, no stimulus was presented in 20 pseudorandomized trials (catch trials) out of 120 trials in every experimental block. In total, participants completed 7 blocks. Each block lasted for ∼10 mins. After each block, stimulation intensities were readjusted to a threshold level.

**Figure 1.**
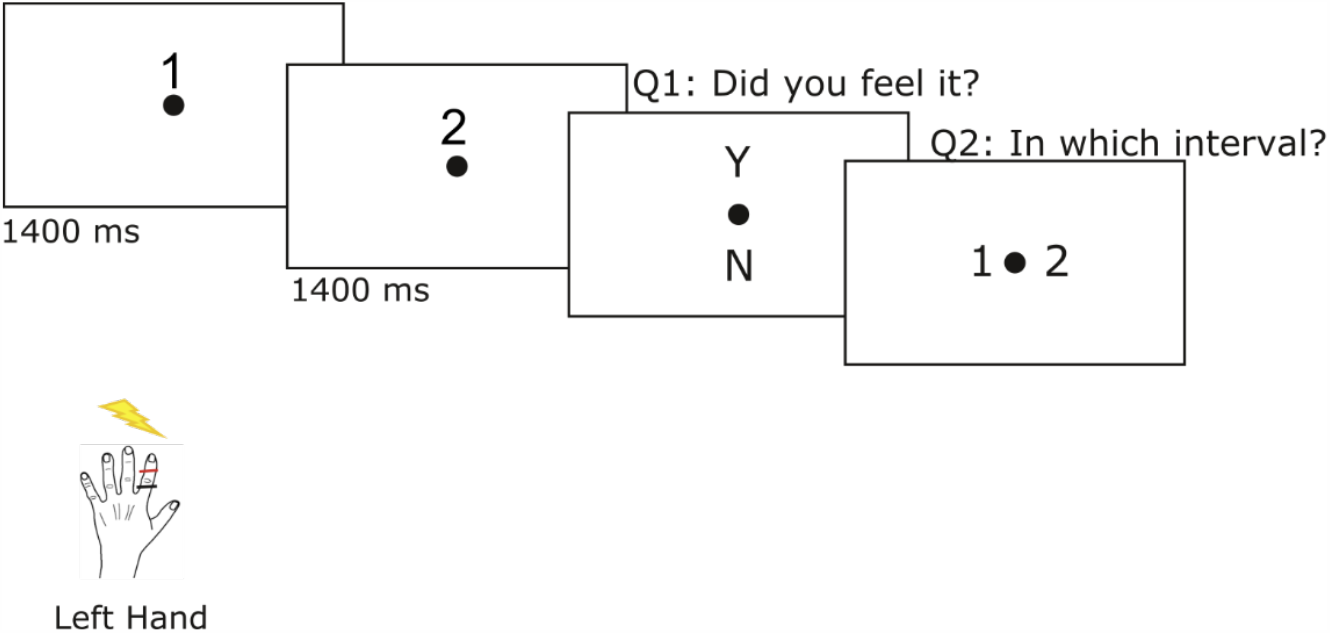
Experimental paradigm. Thirty-six participants received a weak electrical pulse to the left index finger in 720 out of 840 trials over seven experimental blocks. Participants were told that one of the two temporal windows in every trial contained a stimulus; however, no stimulus was actually presented in 120 pseudorandomized trials. In every trial, participants first performed a yes/no detection task and then a two-alternative forced choice task.

### 2.4 Data and code availability

Data and code will be available upon request. Due to a lack of consent of the participants, structural MRI and EEG data cannot be shared publicly and can only be shared upon request if data privacy can be guaranteed according to the rules of the European General Data Protection Regulation (EU GDPR).

### 2.5 Cardiac Cycle Analyses

The changes in somatosensory detection were evaluated across the two phases of the cardiac activity: systole and diastole. Systole was operationalized as ‘the time between the R-peak and the end of the t-wave’ (Motyka et al., 2019; Al et al., 2020). This systolic window length was then used to define a ‘diastolic window of equal length at the end of each cardiac cycle’ (Al et al., 2020). By defining equal lengths of systole and diastole, we balanced the probability of stimulation during both phases (Al et al., 2020). For each cardiac cycle, we used a trapez area algorithm to determine the end of the t-wave (Al et al., 2020; Motyka et al., 2019; Vázquez-Seisdedos, Neto, Marañón Reyes, Klautau, & Limão de Oliveira, 2011). As a result, the average systole (and diastole) length was 338 ± 24ms.

### 2.6 EEG and ECG Recordings

EEG data were recorded from 62 scalp electrodes distributed according to the international 10– 10 system, using a commercial EEG acquisition system (actiCap, BrainAmp; Brain Products) with a bandwidth of 0.015-1000 Hz and a sampling rate of 2500 Hz. During the measurement, the EEG signal was referenced to FCz and an electrode on the sternum served as ground. Electrode impedance was kept ≤5 kΩ for all channels. The position of the individual electrodes was measured in 3D space for each subject using a Patriot motion tracker (Polhemus). To record heart activity, an ECG electrode was mounted under the left breast of the subjects. EEG and ECG data were first acquired during a 2×4 mins eyes-closed resting-state condition and then during the somatosensory task.

### 2.7 Data Preprocessing

EEG data were preprocessed using EEGLAB (Delorme & Makeig, 2004) and custom-written scripts in MATLAB (MathWorks Inc.). A low-pass finite impulse response filter (112.5 Hz) was applied before downsampling EEG data to 250 Hz. All experimental blocks were concatenated and then passed through first a high- and then a low-pass filter (0.3 – 45 Hz) using the fourth order Butterworth filter. The removal of the noisy EEG channels with a flat line longer than 5s or correlating less than 85% with its reconstructed activity from the rest of the channels was followed by their interpolation from their neighboring channels (using “clean_rawdata” plug-in of EEGLAB). After application of principal component analysis, an independent component analysis (ICA) was performed using an extended infomax algorithm to remove the ocular and heartbeat artifacts (Delorme, Palmer, Onton, Oostenveld, & Makeig, 2012). To specifically remove the heartbeat-related artefactual ICA components, the ECG R-peak positions were first identified using the HEPLAB toolbox, which was followed by a visual correction (Perakakis, 2019). Then, ICA data were copied and epoched according to R-peak positions (−100 to 800 ms) to ease the selection of the heartbeat artifacts that show a similar time-course with the R-peak and t-wave of the ECG activity. Artefactual independent components were then visualized using the SASICA software (Chaumon, Bishop, & Busch, 2015) and selected manually for rejection. After removal of the artefactual ICA components, the continuous artifact-free components were forward projected. EEG data were then rereferenced to an average reference.

### 2.8 Somatosensory-Evoked Potential (SEP) Analyses

EEG data were epoched between −1000 to 2000 ms around the stimulus onset individually for trials with stimulation during systole and diastole. A baseline correction was done using the − 100 to 0ms window. The calculation of ‘maximum positive deflection of the early SEP component P50 (40 to 60 ms)’ (Al et al., 2020; Nierhaus et al., 2015; Zhang & Ding, 2010) suggested the C6 electrode as the representative electrode of the contralateral primary somatosensory area. Since the representative electrode was determined as C4 in our previous study (Al et al., 2020), we selected a cluster of electrodes neighboring the C6 electrode (C4, CP4, C6, CP6) broadly covering brain structures relating to somatosensory processing. Statistical analyses of SEP amplitudes were performed on these 4 electrodes. Following our previous study, we also estimated and cleaned artefactual effects of blood circulation on the EEG data (see Al et al., 2020 for details of the ECG-induced artifact cleaning on the SEPs).

### 2.9 Heartbeat-Evoked Potential (HEP) Analyses

In our analysis of HEP, we selected the trials where the somatosensory stimulation occurred ‘at least 400ms after the preceding R-peak (corresponding to diastole)’ to ensure that the time window for neural responses to heartbeats was not contaminated by stimulation-related activity. After this trial selection, we determined HEPs by segmenting data around the R-peak which allowed us to evaluate prestimulus HEPs, often reported between 250 to 400 ms after the R-peak (Kern, Aertsen, Schulze-Bonhage, & Ball, 2013; Schandry, Sparrer, & Weitkunat, 1986; Schandry & Weitkunat, 1990).

### 2.10 Source Reconstruction Analyses

The neural sources of the EEG signal were reconstructed on the BrainStorm toolbox (Tadel, Baillet, Mosher, Pantazis, & Leahy, 2011) using individually measured electrode positions (in 29 subjects). Standard electrode positions were used for 7 subjects since some problems were identified in their recorded electrode positions. When available, individual brain anatomies (17 subjects) and otherwise a template brain anatomy (ICBM152; (Fonov, Evans, McKinstry, Almli, & Collins, 2009) were used. The structural T1-weighted MRI images were segmented using Freesurfer (http://surfer.nmr.mgh.harvard.edu/) and a 3-shell boundary element model (BEM) was constructed to compute the lead field matrix with OpenMEEG (Gramfort, Papadopoulo, Olivi, & Clerc, 2010; Kybic et al., 2005). Then, the lead field matrices were inverted using eLORETA separately for each condition and subject. The Matlab code for eLORETA is derived from the MEG/EEG Toolbox of Hamburg (METH). Individual source data were then baseline-normalized with −100 to 0 ms time-window and then projected to the ICBM152 template. The segmentation of the cortical anatomy was performed according to the Destrieux atlas (Destrieux, Fischl, Dale, & Halgren, 2010).

### 2.11 Signal Detection Theory (SDT) Analyses

In the presence of a stimulus, a trial was categorized as a hit if subjects reported detecting its presence and as a miss if they reported not feeling it. A false alarm (FA) occurred when subjects reported detecting the presence of a stimulus in its absence in the yes/no detection task. Using the hit and FA rates, we calculated signal detection theory measures to differentiate sensitivity (d’) and criterion effects in the yes/no detection task (Al et al., 2020). In our SDT analysis, we determined the position of FAs in the cardiac cycle depending on the expected stimulation timing in the middle of the temporal intervals that subjects allocated the FAs into.

### 2.12 Statistical Analyses

As we described in our previous study, we statistically tested the two-condition comparisons of EEG data by using cluster-based permutation t-tests in the FieldTrip toolbox (Oostenveld, Fries, Maris, & Schoffelen, 2011). For comparison of the conditions regarding the sensory-level SEP activity, cluster statistics were done over a set of somatosensory electrodes (C4, CP4, C6, CP6) either for the mean activity across the previously identified time window of 268 – 468 ms or for the whole time window of 0 – 600 ms. For comparison of HEP activity preceding hits and misses, we tested mean differences between 296 – 400 ms over a cluster of electrodes (FC2, Cz, C4, CP1, CP2, Pz, P4, C1, C2, CPz, CP4, P1, P2) according to our previous study (Al et al., 2020). This analysis was also performed for comparing HEP differences during the resting state and task.

For the statistical test of the neural sources of differential HEP amplitudes, we used cluster statistics to compare mean HEP amplitude between 296 – 400 ms following the R-peak between hits and misses in a dataset of 37 subjects from our previous study (Al et al., 2020) over 15,000 cortical vertices. The resulting cortical regions were further used as the region of interest for the HEP analyses in the present dataset.

For the within-subject ANOVA tests, we checked for the sphericity assumption and if it was violated, then we corrected results by Greenhouse–Geisser method. All statistical tests were two-sided.

## 3. Results

Thirty-six participants were presented weak somatosensory (electrical) stimuli to the left index finger in a combined yes/no and 2IFC detection task (**Fig. 1**). Both EEG and electrocardiography (ECG) were recorded. On average, participants detected 52.2± 10.0% (mean ± SD) of the somatosensory stimuli with a false alarm rate of 8.6 ± 11.4%. Participants correctly detected 79.4 ± 5.6% of the stimulation interval.

### 3.1 Somatosensory Perception Changes across the Cardiac Cycle

We first tested our hypothesis that somatosensory detection is less likely during systole compared to diastole phase of the cardiac cycle as we found in our previous studies (Al et al., 2020; Motyka et al., 2019). For this purpose, we determined when stimulus onsets occurred during systole and diastole. As we expected, somatosensory detection was significantly lower during systole (M= 50.37%) than diastole (M= 53.22%), *t*_35_ = −3.41, *p* = 2·10^−3^ (**Fig. 2a**). 27 out of 36 participants had a lower detection rate during systole. We furthermore checked whether this cardiac phase effect on detection was correlated with individual differences in the heart rate or the heart rate variability (HRV, i.e., the standard deviation of RR intervals). We expected to find a significant correlation with HRV depending on our previous results (Al et al., 2020). However, subject’s detection rate difference between the two cardiac phases did not significantly correlate either with their heart rate or HRV (respectively, Pearson’s correlation, *r* = −0.05, *p* = 0.77 and *r* = 0.21, *p* = 0.22).

**Figure 2.**
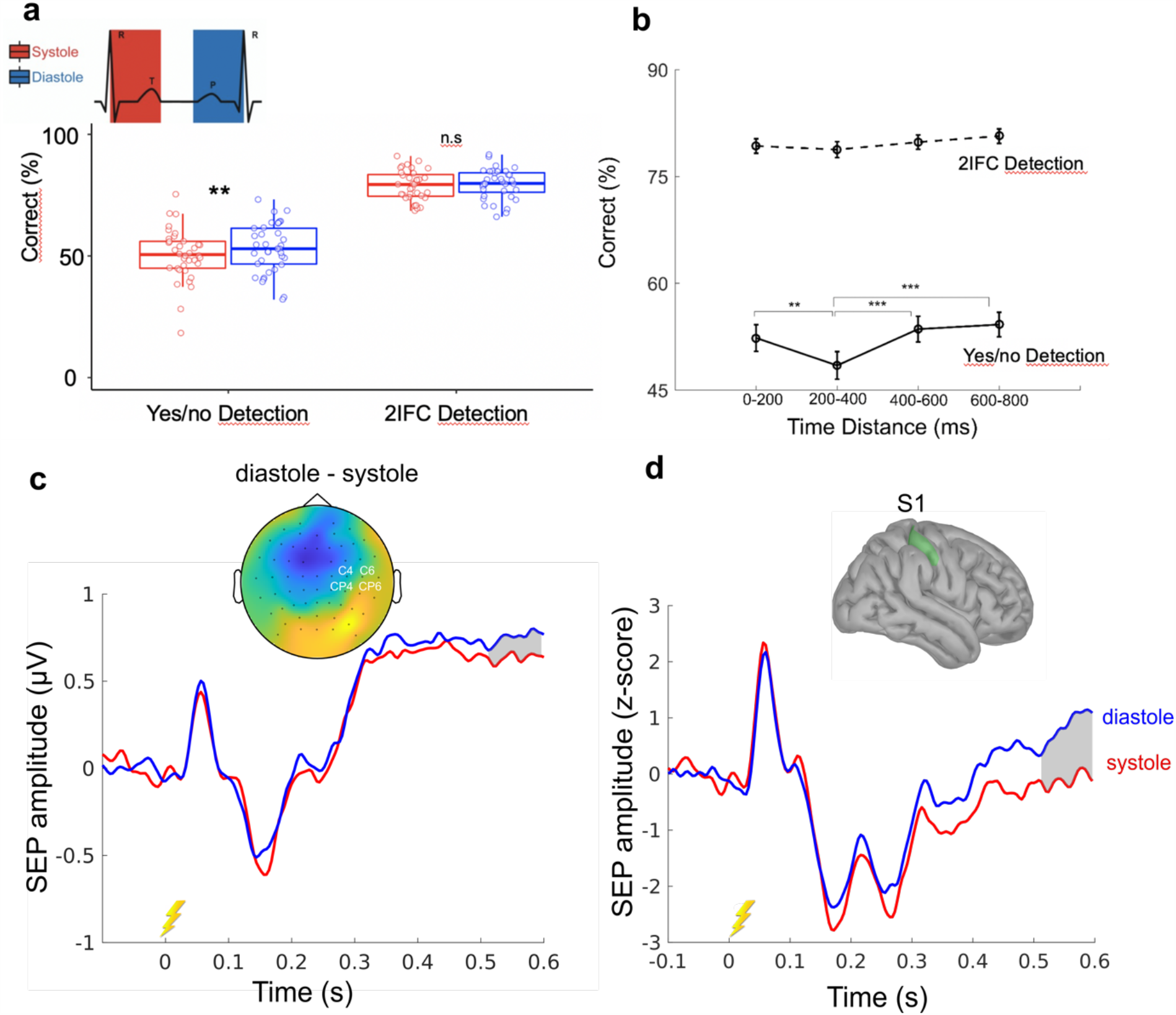
Somatosensory perception and evoked potentials across the cardiac cycle. (a) Correct detection of stimulus presence (yes/no) and its temporal interval (2IFC) during systole and diastole phase of the cardiac cycle. When a stimulus coincided with systole compared to diastole, participants detected its presence less often (*t*_35_ = − 3.35, *p* = 2·10^−3^). However, their ability to correctly identify the stimulation interval did not significantly change between systole and diastole (*p* = 0.28). (b) Correct yes/no and 2IFC detection of somatosensory stimuli relative to their distance from the previous R-peak. Yes/no detection performance was lowest 200 – 400 ms after the R-peak. (*post-hoc* paired *t-*test between 200 – 400 and 600 – 800 ms, *t*_35_ = − 4.22, *p* = 2·10^−4^) (c) Grand average somatosensory-evoked potentials (SEPs) for electrical stimulations during systole and diastole. The P300 component of SEPs was less positive for stimuli during systole than diastole between 512 – 600 ms after stimulus onset at a cluster of electrodes (indicated with the white text) around the contralateral somatosensory cortex (Monte Carlo *p* = 0.004). The topography plot shows the contrast between diastole and systole between 512 and 600 ms. The source-reconstructed P300 amplitude was significantly different between systole and diastole in the contralateral somatosensory cortex (S1) similar to the sensory data (*t*_35_ = −2.78, *p* = 0.01). Error bars represent SEMs. \**p* < 0.05, \*\**p* < 0.005, \*\*\**p* < 0.0005, ns, not significant.

We additionally tested the effect of stimulation interval and cardiac phase on detection rates (**Supp. Fig. 1**). A within-subject ANOVA test confirmed the increased detection rates during diastole (*F*_1,35_ = 11.42, *p* = 2·10^−3^) and showed that stimuli during the second temporal interval were detected more often (*F*_1,35_ = 27.60, *p* = 7·10^−6^). We did not observe a significant interaction effect between the temporal interval and cardiac phase (*F*_1,35_ = 0.99, *p* = 0.3).

Next, we tested whether the changes in the yes/no detection rates were associated with changes in perceptual sensitivity (*d’*) according to Signal Detection Theory (SDT). Confirming our previous finding, detection sensitivity was found to be significantly lower during systole (M= 1.43) relative to diastole (M= 1.75), *t*_35_ = −3.77, *p* = 6·10^−4^ (**Supp. Fig. 2a**). For the criterion, we observed a trend between systole (M= 0.71) and diastole (M= 0.79), *t*_35_ = −1.95, *p* = 0.06 (**Supp. Fig. 2b**). Furthermore, the effect of the cardiac phase on 2IFC detection was tested. No significant differences were observed in 2IFC detection between systole (M= 78.91%) and diastole (M= 79.54%), *t*_35_ = −1.13 *p* = 0.27 (**Fig. 2a**).

Somatosensory detection was previously shown to be minimum when stimulation occurred during 200 – 400 ms after the previous heartbeat (Al et al., 2020). To test the reliability of this result, we tested the effect of the time delay of somatosensory stimulation from the previous R-peak on yes/no detection performances. Within-subject ANOVAs confirmed that detection performance significantly differed between four time windows: 0 – 200, 200 – 400, 400 – 600, and 600 – 800 ms (*F*_3,105_ = 9.58, *p* = 1·10^−5^). Also, a trend for 2IFC detection was observed (*F*_3,105_ = 2.36, *p* = 0.07). Yes/no detection was lowest 200 – 400 ms after the R-peak (*post-hoc* paired *t-*test between 0 – 200 and 200 – 400 ms windows for yes/no detection: *t*_35_ = 3.32, *p* = 0.002; between 200 – 400 and 600 – 800 ms for yes/no detection: *t*_35_ = − 4.22, *p* = 2·10^−4^, **Fig. 2**).

### 3.2 Somatosensory-Evoked Potentials Change across the Cardiac Cycle

Similar to perceptual attenuation, the amplitude of a late component of somatosensory-evoked potentials (SEPs), P300, has been also previously reported to decrease during systole. Therefore, we expected to observe a decreased P300 activity in response to stimulation during systole compared to diastole. To test this hypothesis, we first checked the mean P300 difference in the time window of 268 – 468 ms, as previously reported, between systole and diastole over the contralateral somatosensory cortex (indexed by C4, C6, CP4, CP6). The cluster statistics testing the mean SEPs between 268 – 468ms over the somatosensory electrodes showed only a trend effect, suggesting a lower mean P300 activity during systole compared to diastole (Monte Carlo *p* = 0.07, corrected for multiple comparisons in space). Then, we performed a cluster-based permutation *t*-test comparing SEPs during the two cardiac phases between 0 – 600 ms following stimulus onsets over the contralateral somatosensory cortex. The results showed that during a later time window, 512 – 600 ms following stimulation during systole, late SEPs showed decreased positivity (Monte Carlo *p* = 0.02 respectively, corrected for multiple comparisons in time and the region of interest electrodes; **Fig. 3a**). To confirm that the SEP differences across the cardiac cycle originate from the contralateral somatosensory cortex (S1), we also performed a source reconstruction analysis (**Fig. 2**). Source analysis confirmed that late P300 amplitude in the S1 was significantly different between the two cardiac phases (*t*_35_ = − 2.78, *p* = 0.01; **Fig 2**).

**Figure 3.**
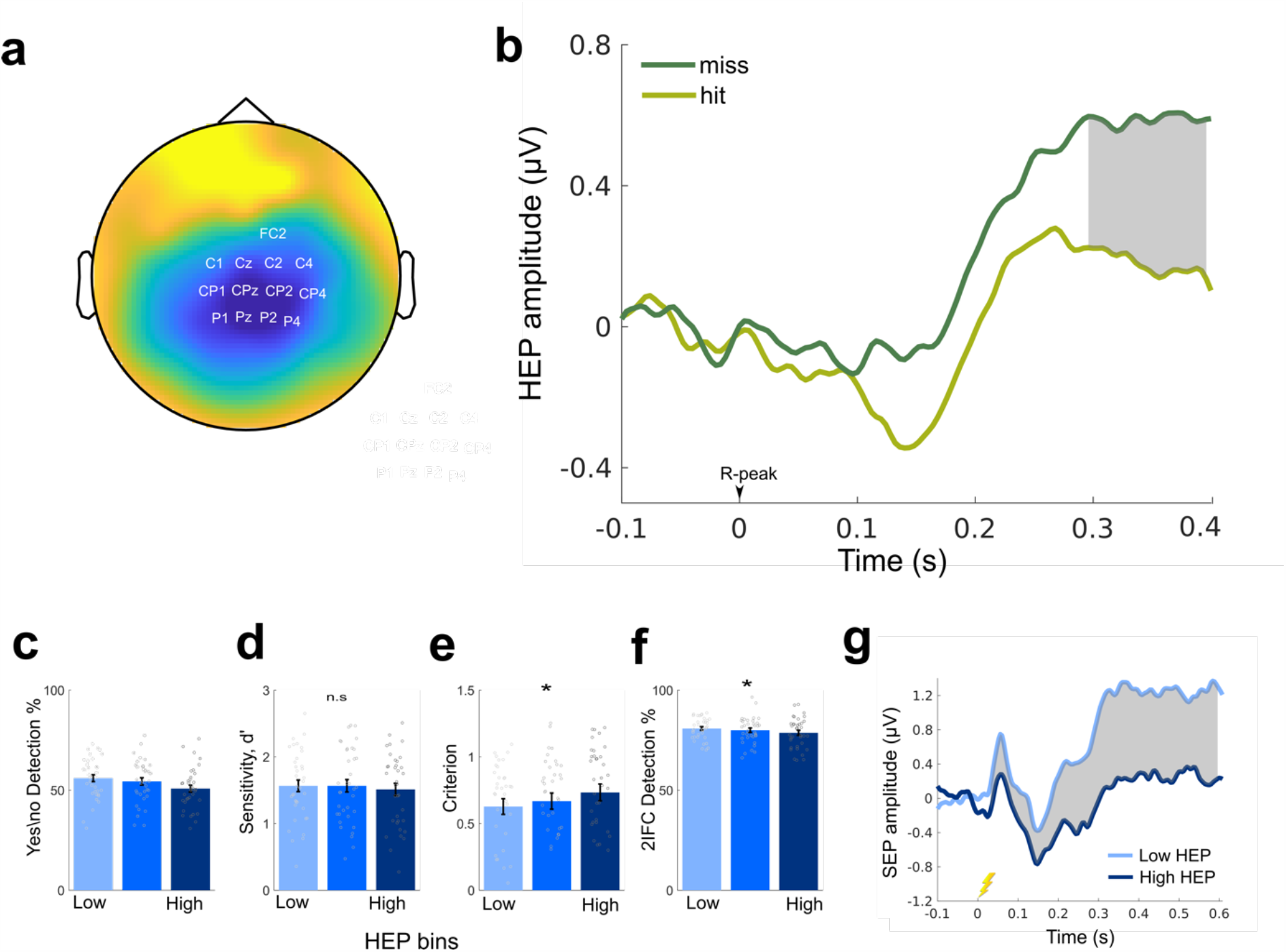
The effect of prestimulus HEPs on the upcoming somatosensory perception. (a) The topography contrast of prestimulus HEP amplitude preceding hits and misses between 296 – 400ms following the R-peak. The region of interest electrodes that were used in the HEP analyses is highlighted. (b) In 36 participants, mean HEP amplitudes between 296 – 400 ms were higher preceding misses than hits across the central electrodes (Monte Carlo *p* = 0.001) (c-f) For every single trial, the mean HEP amplitude was calculated across the indicated electrodes in the 296 to 400 ms time window. Then, single trials were sorted depending on the mean HEP amplitude and divided into three equal bins for each participant. (c) Yes/no detection rate decreased as HEP amplitude increased. (d) This change in detection of stimulus presence was not correlated with a significant decrease in sensitivity (*p* = 0.6), (e) however connected with an increase in criterion, i.e., having a more conservative decision bias and therefore reporting the absence of stimulus more often (*p* = 0.04). (f) Similarly, the correct allocation of stimulus into its temporal interval decreased as HEP amplitudes increased (*p* = 0.03). The individual participant data are shown on the bar plots as gray points. (g) SEP amplitudes following low and high HEP amplitudes. After the low compared high HEP amplitudes, mean SEPs between 32 and 600 ms following stimulation were significantly more positive over contralateral somatosensory electrodes (C4, CP4, C6, CP6; Monte Carlo P = 0.001). Error bars represent SEMs. **P < 0.005, ***P < 0.0005; ns, not significant.

We additionally checked SEP amplitudes separately for stimulations presented in the 1^st^ and 2^nd^ temporal intervals during systole and diastole over S1 in the source level. (**Supp. Fig. 3**). While mean late P300 amplitudes were significantly lower during systole compared to diastole for stimulations during the 1^st^ interval (*t*_35_ = −2.70, *p* = 0.01), no significant effect of cardiac phase on SEPs was observed for stimulations during the 2^nd^ interval (*t*_35_ = −1.41 *p* = 0.17).

### 3.3 Heartbeat-Evoked Potentials Negatively Correlate with Upcoming Somatosensory Detection and Evoked-Potentials

Following our previous study, we hypothesized that HEPs preceding stimulus onset predicts somatosensory detection. More specifically, we expected to observe higher levels of HEP amplitudes preceding misses compared to hits across the central electrodes (FC2, Cz, C4, CP1, CP2, Pz, P4, C1, C2, CPz, CP4, P1, P2) as reported by Al et al. (2020). In the present study, prestimulus mean HEPs between 296 – 400 ms following the R-peak were again found to be higher before misses than hits over the central electrodes (Monte Carlo *p* = 0.001, corrected for multiple comparisons across the region of interest electrodes; **Fig. 3a**). We furthermore confirmed the prestimulus HEP differences between hits and misses separately for stimulations during the first and second temporal intervals (**Supp. Fig. 4**)

Because of our previous results, we expected this HEP difference to be reflected in a change in criterion (detection bias) rather than sensitivity according to SDT. To test this hypothesis, we sorted single trials according to mean HEP amplitude (across the central electrodes in the 296 – 400 ms time window) and split them into three equal bins for each participant. Again, we found that detection rates decreased with increasing HEP amplitudes (To avoid double-dipping, we did not apply a statistical test here). This decrease in detection was paralleled with an increase in detection criterion (within-subject ANOVA, *F*_2, 70_ = 3.37, *p* = 0.04, **Fig. 3e**). In other words, when HEP levels were higher, participants were more conservative to report that they felt a stimulus. No significant changes were observed in sensitivity over HEP bins (*F*_2,70_ = 0.39, *p* = 0.6, **Fig. 3d**). We then tested whether prestimulus HEP amplitude also affected subject’s ability to correctly detect the temporal interval of the stimulation. Similar to yes/no detection, 2IFC detection decreased as HEP levels increased (*F*_2,70_ = 3.59, *p* = 0.03, **Fig. 3f**).

We furthermore tested the effect of prestimulus HEP amplitudes on the upcoming SEP amplitudes. Given our previous results, we expected that following lower HEP amplitudes, the upcoming SEPs in the time window of 32 – 600 ms (0= stimulus onset) had higher positivity. Confirming our previous findings, following low compared to high HEP amplitudes, we observed higher positivity for SEPs between 32 and 600 ms after stimulation over the somatosensory electrodes (Monte-Carlo *p* = 0.001; **Fig. 3g**). The source analysis also confirmed that following low and high HEP levels, the amplitude of the early SEP component, P50, was significantly different in the S1 (*t*_35_ = 2.40, *p* = 0.02). As we observed this P50 modulation also in other brain areas known to play a role in heart-brain interactions such as the right anterior insula, the left and right posterior cingulate cortex (PCC), and the left and right lateral prefrontal cortex (LPFC) in our previous study, we again tested P50 differences in those areas. Similarly, following low and high HEP levels, P50 amplitudes were found to be significantly different (after false discovery rate correction, significant p<0.01) in the right anterior insula (*t*_35_ = 2.98, *p* = 0.005), the left and right PCC (*t*_35_ = −5.77 *p* = 2·10^−6^ and *t*_35_ = − 6.23, *p* = 4·10^−7^), and the left LPFC (*t*_35_ = −2.77, *p* = 0.01). Only a trend was observed in the right LPFC (*t*_35_ = −2.63, *p* = 0.01).

In an exploratory analysis, we also tested overall changes in HEP levels preceding hits and misses across all electrodes for the entire window of HEP, *250 – 400 ms after the R-peak*. A cluster-based permutation *t-*test revealed a significant positive cluster over frontal electrodes and a negative cluster over central electrodes between 250 – 400 ms (respectively, Monte-Carlo *p* = 0.001 and *p* = 0.001 corrected for multiple comparisons in space and time; **Supp. Fig. 5)**. The significant result including a negative cluster over central electrodes replicated our hypothesis-driven analyses, where misses were preceded by higher positivity of HEP. The frontal regions in the positive cluster had a negative HEP polarity and the absolute strength of prestimulus HEP for misses was similarly higher than hits. Since there were no significant differences in ECG amplitude preceding hits and misses (no clusters were found), the HEP differences in neural data cannot be explained by volume conduction of cardiac electrical activity.

### 3.4 Neural Sources of the Differential Heartbeat-Evoked Potentials

To reveal the cortical regions contributing to the differential responses to prestimulus heartbeats in hits and misses, we first ran source reconstruction analysis on the EEG dataset from our previous study (Al et al., 2020). The cluster statistics contrasting mean HEP amplitudes between 296 – 400 ms across all cortical vertices between hits and misses showed that two regions in the right hemisphere were differentially activated: one extending from the postcentral gyrus and sulcus, the paracentral lobule and sulcus to the superior parietal lobule (Monte Carlo *p* = 0.02, **Fig. 4a**) and another extending across the precuneus, isthmus cingulate, middle and posterior cingulate cortex, pericallosal sulcus (Monte Carlo *p* = 0.02, **Fig. 4b**). Then, we used these two regions as our region of interest to test mean HEP differences preceding hits and misses in our recent study. Our analyses confirmed the differential mean HEP activity between 296 – 400 ms in both regions. (the parietal areas: *t*_35_ = − 2.44, *p* = 0.02, **Fig. 4c** and the cingulate areas: *t*_35_ = 2.86, *p* = 0.01, **Fig. 4d**).

**Figure 4.**
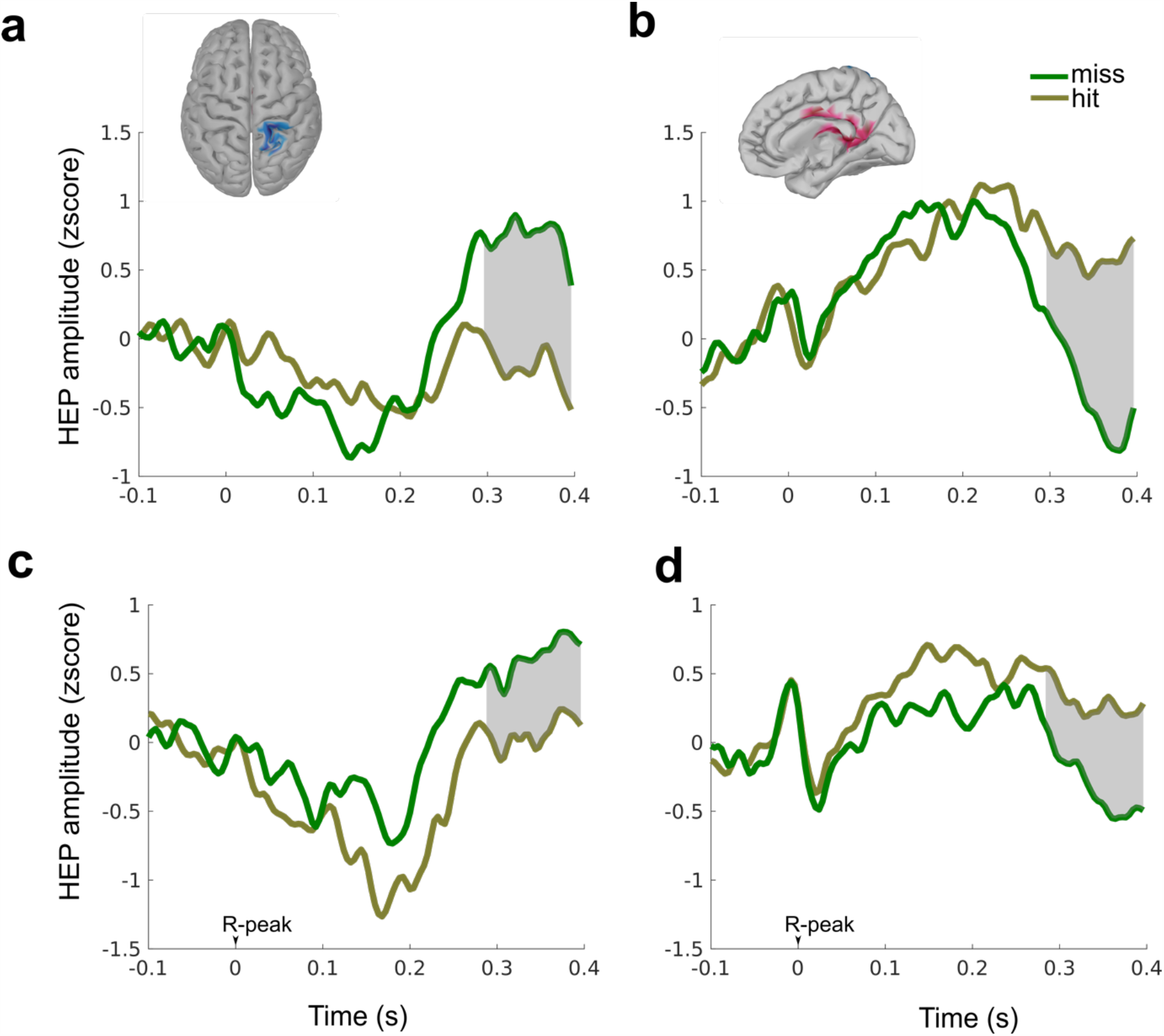
Neural sources of the differential heartbeat-evoked potentials (HEPs) preceding detection. (**a-b**) The neural sources of HEP differences preceding hits and misses were determined in a previous EEG dataset with 37 subjects from the study of Al et al. (2020). Cluster statistics comparing mean HEP amplitudes between 296 – 400 ms following the R-peak across all cortical vertices for hits and misses revealed two clusters: (**a**) one extending from the postcentral gyrus and sulcus, the paracentral lobule and sulcus to the superior parietal lobule (Monte Carlo *p* = 0.02 corrected for multiple comparisons in space and time) and (**b**) another extending across the precuneus, isthmus cingulate, middle and posterior cingulate cortex, pericallosal sulcus (Monte Carlo *p* = 0.02). (**c-d**) In the present study, mean HEP amplitudes (between 296 – 400 ms) preceding hits and misses were significantly different over (**c**) the parietal regions (indicated with blue in the cortex, *t*_35_ = − 2.44, *p* = 0.02 and (**d**) the posterior cingulate areas (indicated as red, *t*_35_= 2.86, *p* = 0.01), confirming HEP differences from the first dataset.

### 3.5 Heartbeat-evoked potentials decrease during a somatosensory task compared to a resting-state condition

If HEP amplitudes decrease as a result of externally oriented attention, then HEP levels are expected to be lower during a somatosensory task compared to a resting-state condition. To test this hypothesis, we contrasted HEP amplitudes from a resting-state condition with HEP amplitudes preceding stimulus during the somatosensory task. Cluster statistics comparing mean HEP amplitudes between 296 – 400 ms following the R-peak confirmed that HEP amplitudes were lower during the task compared to a resting-state condition over the central electrodes (Monte-Carlo *p* = 0.01 corrected for multiple comparisons in space, **Fig. 5a**). We furthermore compared heart rate during the resting state and the task. No significant differences in heart rate were found (*t*_35_ = 0.63, *p* = 0.53).

**Figure 5.**
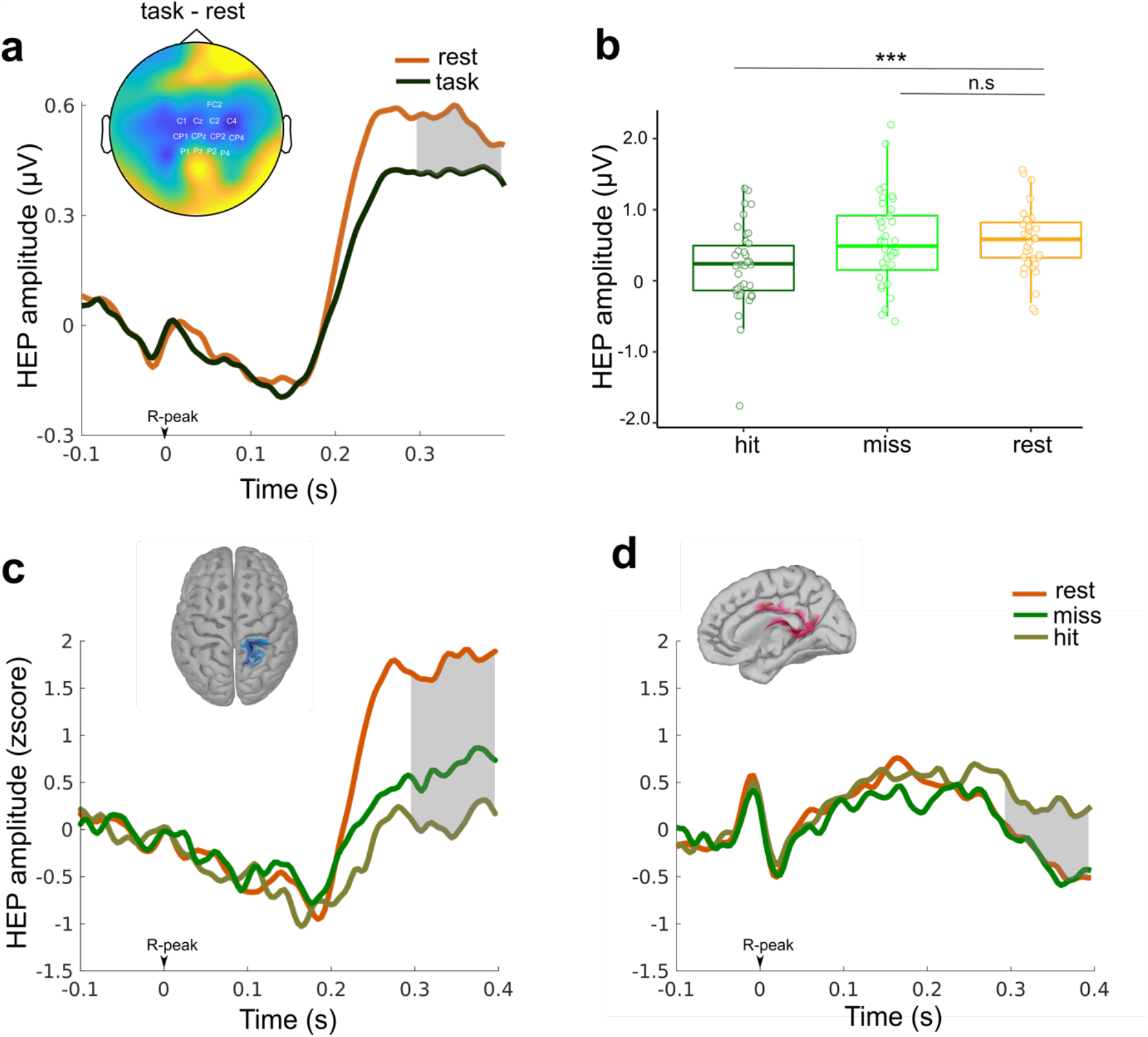
HEP amplitudes during a somatosensory task compared to a resting-state condition. (**a**) HEP amplitudes over a cluster of central electrodes (indicated with the white text on the topoplot) between 296 – 400 ms were lower during the task compared to a resting-state condition (Monte-Carlo *p* = 0.01 corrected for multiple comparisons in space). (**b**) We then compared mean HEP amplitudes (across the indicated cluster electrodes between 296 – 400 ms) during the resting state with HEP amplitudes preceding hits and misses. During the resting-state condition, HEP amplitudes were higher than those preceding hits (*t*_35_ = 4.12, *p* = 2·10^−4^) but not misses (*t*_35_ = −0.04, *p* = 0.97). (**c**) Source-reconstructed HEP activity during a resting-state condition and before detecting or missing a somatosensory stimulus are shown in the parietal regions extending from the postcentral gyrus and sulcus, the paracentral lobule and sulcus to the superior parietal lobule. In the parietal areas, HEPs during the resting-state were significantly different from those preceding hits (*t*_35_= 3.03, *p* = 4·10^−3^) and showed a trend against misses (*t*_35_ = 2.03, *p* = 0.05). (**d**) In the predefined regions around cingulate areas (indicated with red), HEP amplitudes during the rest were again significantly different than those preceding hits (*t*_35_ = −2.13, *p* = 0.04). In this region, no significant differences were found between resting-state HEPs in comparison to those before misses (*t*_35_ = 0.11, *p* = 0.91).

We then compared mean HEP amplitudes during the resting state with those preceding hits and misses (indicated in **Fig. 5b**). HEPs during rest were higher than those preceding hits (*t*_35_ = 4.12, *p* = 2·10^−4^) but not misses (*t*_35_ = −0.04, *p* = 0.97). Thereafter, we contrasted source-reconstructed HEP amplitudes during rest with those preceding hits and misses over the parietal and cingulate areas as these regions were determined in the previous section (see *3.4* for details). In the parietal areas, mean HEPs between 296 – 400 ms during the resting-state were significantly different from those preceding hits (*t*_35_ = 3.03, *p* = 4·10^−3^) and showed a trend against misses (*t*_35_ = 2.03, *p* = 0.05; **Fig. 5c**). In the posterior cingulate areas, mean HEP amplitudes during the rest were again significantly different than those preceding hits (*t*_35_ = −2.13, *p* = 0.04) but not misses (*t*_35_ = 0.11, *p* = 0.91; **Fig. 5d**).

## 4. Discussion

The findings of our study replicate the effect of stimulus timing during the cardiac cycle and prestimulus HEP amplitudes on somatosensory perception and evoked potentials. Somatosensory detection is again found to be lower during systole compared to diastole and to correlate negatively with prestimulus HEP amplitudes. Consistently, the cardiac phase effect on detection is associated with changes in detection sensitivity whereas the HEP effect correlated with criterion. Moreover, the cardiac phase is again found to affect only late SEP components while prestimulus HEP amplitudes influence both early and later components. The new source analyses on two different datasets shows that HEP modulations specifically in the right posterior cingulate and parietal areas relate to changes in somatosensory perception. We furthermore show that subjects’ HEP amplitudes preceding misses during the somatosensory task approached their level during the resting-state, when exteroceptive attention is expected to reach a minimum.

We previously interpreted the changes in the detection and neural responses across the cardiac cycle in an interoceptive predictive coding framework. In this framework, rhythmic bodily fluctuations such as the heartbeat are treated as predictable events and the resulting physiological changes in the body are suppressed from being consciously perceived. In this way, the brain can minimize interference from self-generated signals such as heartbeat-related firing changes (Barrett & Simmons, 2015; Seth & Friston, 2016). For example, afferent neurons in the fingers have been shown to fire in response to the changes in the pressure wave, which can be predicted and suppressed by the central processes (Macefield, 2003). During this phase, it is likely that the perception of a weak somatosensory stimulus could be also suppressed with other heartbeat-related signals. In our recent study, stimulus detection was again observed to be minimal when stimuli were presented during 200 – 400 ms following the heartbeat (R-peak), roughly coinciding with the pulse-wave. Therefore, our new findings support the predictive mechanisms as described above. By measuring the finger pulse directly, future studies might investigate precise timing of this suppression related to features of the pulse-wave, i.e., initial arrival, steepest rise or maximum.

Similar to our previous results, we found that the decrease in yes/no detection rates was reflected in a decrease in detection sensitivity. In other words, during systole, participants were worse at discriminating the presence of the weak somatosensory stimuli from the internal noise. Interestingly, even though we observed significant modulations of yes/no detection rates along the cardiac cycle, we did not observe similar significant changes in the subjects’ ability to allocate a stimulus into a correct temporal interval in the 2IFC detection task. One reason for it might be that participants compared the two temporal intervals corresponding to the different cardiac cycles in the 2IFC task, which might cancel out the cardiac phase effect on the 2IFC detection. Another reason might be related to subjects’ high 2IFC performance compared to their threshold level performance in the yes/no detection task. This might have hindered the effect of cardiac phase on 2IFC performance. A further difference between these two tasks is that while yes/no detection is a subjective task, which is affected by a decision bias, 2IFC detection is an objective, i.e., bias-free, task (Azzopardi & Cowey, 1997). However, since we observed that cardiac phase mainly affected the sensitivity aspect of detection, it is not likely that the objectivity of the 2IFC task hindered the cardiac phase effects on it.

In addition to the decrease of somatosensory perception, the late component of SEPs, P300, was observed to be weaker during systole as compared to diastole. Interestingly, a significant difference in the P300 component was observed between 512 – 600 ms after the stimulation, which was later than the 268 – 468 ms window that was observed in our earlier study (Al et al., 2020). We previously suggested that this decrease in P300 amplitude can be associated with a more accurate prediction of the pulse-wave (as a result of a smaller ‘prediction error’) (Al et al., 2020; Friston, 2005). Despite the different time windows in which the effect was statistically significant, current results support the notion that the weaker P300 amplitude during systole is a result of perceptual suppression of the peripheral activity via central prediction (Al et al., 2020). The smaller P300 amplitude during systole might indicate a less efficient ‘propagation’ of somatosensory information to higher cortical areas (Vugt et al., 2018). Such a decrease in neural propagation could interfere with both the global ‘broadcasting’ of the stimulus (Dehaene, Sergent, & Changeux, 2003) and/or late recurrent activity within somatosensory areas (Auksztulewicz, Spitzer, & Blankenburg, 2012; Lamme, 2006), both of which can suppress the conscious perception of the external stimulus

Not only cardiac phase but also the amplitude of prestimulus HEP affected somatosensory processing. Here we replicated that higher HEP amplitudes between 296 to 400 ms over centroparietal electrodes were followed by decreased somatosensory detection. This decrease in somatosensory detection was related to a more conservative detection bias (criterion). Since a conservative bias is known to relate to lower baseline firing rate in the brain, higher amplitudes of HEP might interfere with upcoming stimulus detection by suppressing the early neuronal activity from reaching the threshold of ‘ignition’ (according to global neuronal workspace theory) (Dehaene & Changeux, 2011; Vugt et al., 2018). In line with this argument, we confirmed the interference of higher HEP amplitudes with both early (P50) and later (N140, P300) SEP amplitudes in the sensory level. After the fluctuations of the HEP amplitude, we furthermore observed significant changes of the source-localized P50 amplitude in the contralateral somatosensory cortex, right insular cortex, lateral prefrontal cortex, and posterior cingulate cortex. Among these regions, the right anterior insula is known to play a crucial role in regulating attention internally and externally, possibly through its connections with the lateral prefrontal cortex and posterior cingulate cortex, which are also associated with attentional control. Since HEP amplitudes have been previously demonstrated to be higher when attention is directed internally compared to externally (García-Cordero et al., 2017; Petzschner et al., 2019; Villena-González et al., 2017), we reiterate our earlier interpretation that increases in the HEP amplitudes preceding misses indicate an shift in attention from external stimuli to internal bodily signals (Al et al.,2020).

Supporting the interpretation of the increased HEP levels as a result of attention being oriented internally, we found that subjects had higher HEP amplitudes when they were resting compared to engaging in a somatosensory task. Furthermore, during the resting state, HEP levels were observed to be specifically lower than those preceding hits but not misses in sensory level. In other words, subjects’ HEP amplitudes preceding misses during the somatosensory task approached their level during the resting-state, when exteroceptive attention is expected to be kept to a minimum. Thus, these results further substantiate that increases in HEP amplitudes reflect a state of mind focused on interoceptive relative to exteroceptive processes, which interferes with conscious somatosensory detection. A specific point why the somatosensory system might be particularly sensitive to modulations of interoception, might be the fact that both interoception and somatosensation rely on processes in the primary somatosensory cortex (Critchley, Wiens, Rotshtein, Öhman, & Dolan, 2004; Khalsa, Rudrauf, Feinstein, & Tranel, 2009; Pollatos, Schandry, Auer, & Kaufmann, 2007).

Source analysis of HEP fluctuations related to detection versus misses of somatosensory stimuli revealed two regional clusters: One extending from the right postcentral gyrus and sulcus, the paracentral lobule and sulcus to the superior parietal lobule and another extending across the right precuneus, isthmus cingulate, middle and posterior cingulate cortex, pericallosal sulcus (see **Fig. 4**). Furthermore, in these regions, HEP levels preceding misses approached the level of those during a resting-state condition. Among these regions, the postcentral gyrus, the paracentral lobule, middle and posterior cingulate cortex, and precuneus have been previously found to show a higher activity during interoception compared to exteroception and to correlate with interoceptive awareness (Stern et al., 2017). Interestingly, a strong functional connection of precuneus and posterior cingulate cortex with sensorimotor areas including the superior parietal lobule, paracentral lobule, and postcentral gyrus has been observed previously (Margulies et al., 2009; Morecraft, Cipolloni, Stilwell-Morecraft, Gedney, & Pandya, 2004; Nierhaus et al., 2015; Vogt & Vogt, 2003). Furthermore, the precuneus and posterior cingulate cortex are important parts of the default mode network (DMN) (Fransson & Marrelec, 2008; Raichle et al., 2001), which shows higher activation during a resting condition relative to external attention-demanding tasks and which has been linked to self-referential mental activity and interoception (Buckner, Andrews-Hanna, & Schacter, 2008; Kleckner et al., 2017). Taken together, these results provide a neural underpinning for the interpretation that higher HEP amplitudes reflect a state of mind focused on internal processes.

In summary, this study confirms the two different effects of cardiac phase and HEP amplitudes on somatosensory perception and evoked potentials. We interpret the cardiac phase effects in an interoceptive predictive coding framework and link the HEP effects with spontaneous switches between interoceptive and exteroceptive attention. Our additional finding that HEP during the task is smaller than during resting state further supports the latter interpretation. Finally, a new source analysis of HEP fluctuation provides a neural underpinning for the switch between interoception and exteroception.

## 5. Acknowledgements

We thank Sylvia Stasch for technical assistance with data acquisition, Eleni Panagoulas for helping with data preprocessing, Dan John Cook for his valuable comments on the manuscript.

## Notes

### Competing Interest Statement

The authors have declared no competing interest.

