## Supplementary Files for "Heartbeat and Somatosensory Perception"

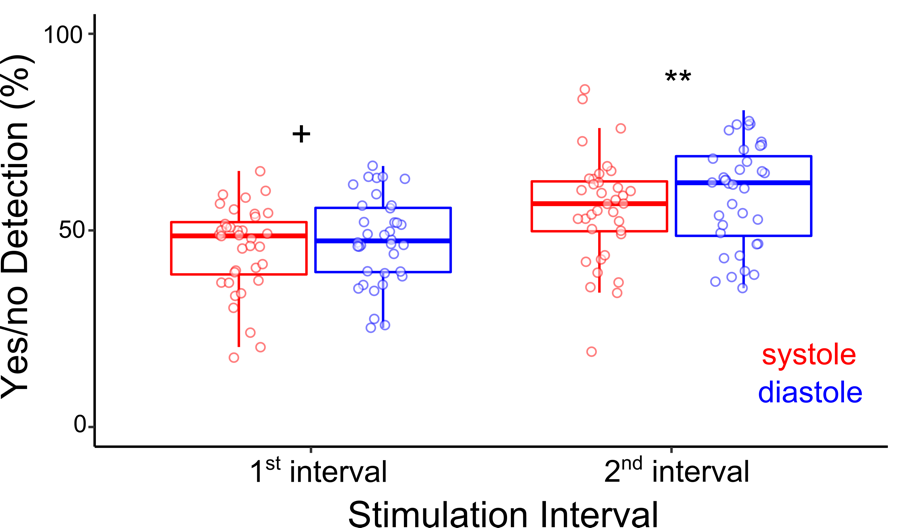

**Supplementary Figure 1**. The effect of stimulation interval and cardiac phase on detection rates. A within-subject ANOVA test confirmed the increased detection rates during diastole (*F*_1,35_=9, *p*=2⋅10^-3^) and showed that stimuli occurred during the second temporal interval were detected more often (*F*_1,35_=27.60, *p*=7⋅10^-6^).

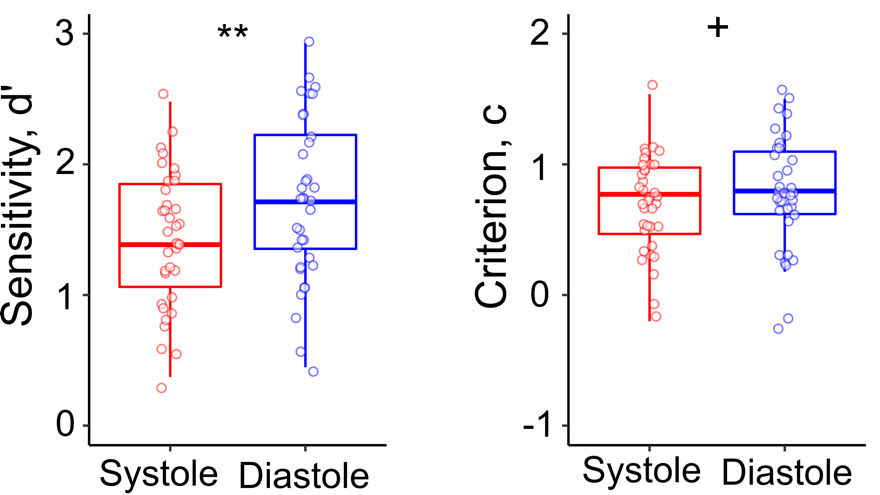

**Supplementary Figure 2.** Detection sensitivity and criterion across the cardiac cycle. (**a**) Subjects had a lower sensitivity to somatosensory stimulation during systole compared to diastole (*t*_35_=-3.77, *p*=6⋅10^-4^). (**b**) Detection criterion showed a trend to be lower during systole (M=0.71) compared to diastole (*t*_35_=-1.94, *p*=0.06).

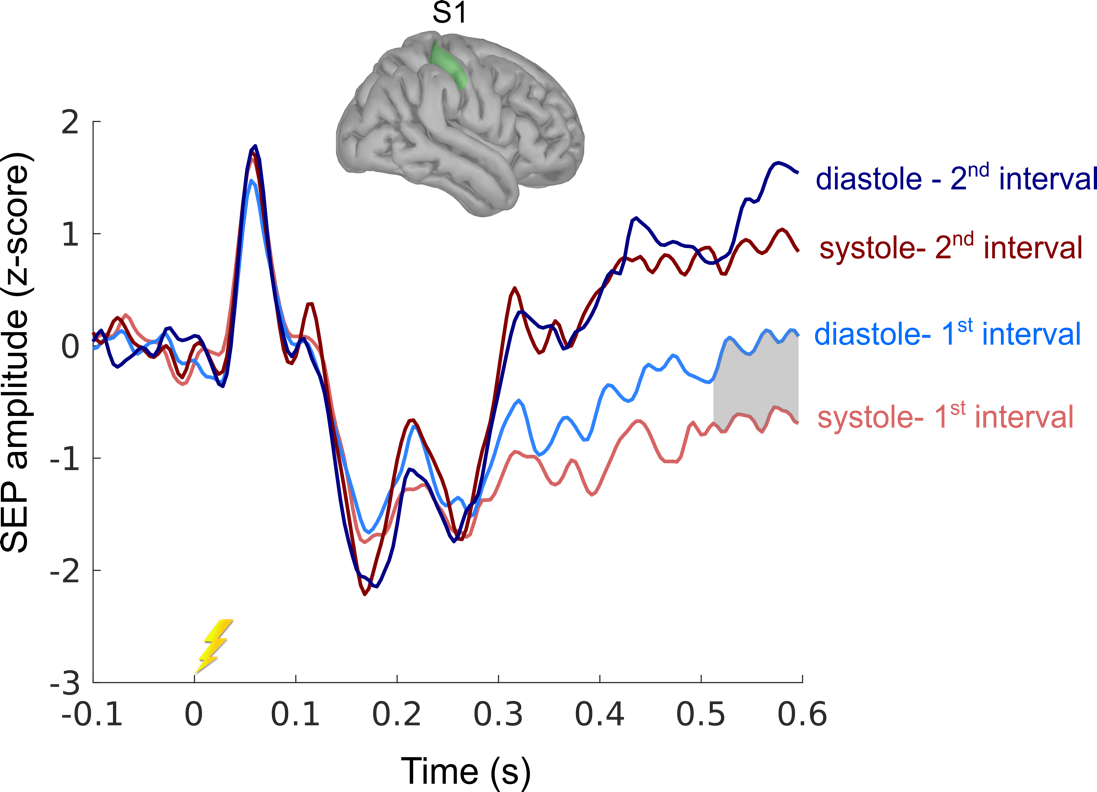

**Supplementary Figure 3.** SEP amplitudes separately for stimulations presented in the 1^st^ and 2^nd^ temporal intervals during systole and diastole in source level. While mean late P300 amplitudes were significantly lower during systole compared to diastole for stimulations during the 1^st^ interval (*t*_35_=-2.70, *p*=0.01), no significant effect of cardiac phase on SEPs was observed for stimulations during the 2^nd^ interval (*t*_35_=-1.41 *p*=0.17).

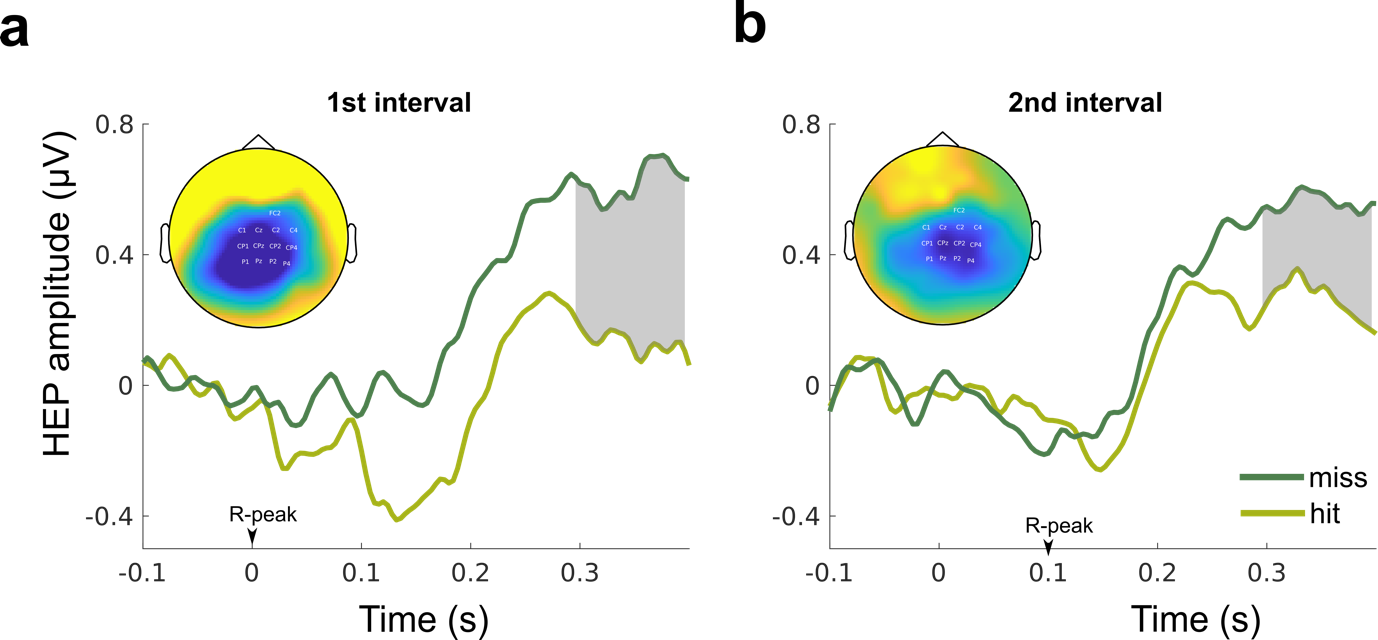

**Supplementary Figure 4.** Prestimulus HEPs for misses and hits during the two temporal intervals. (a) Mean HEP amplitudes between 296–400ms were higher preceding misses compared to hits both when stimulation occurred in (**a**) the first and (**b**) second temporal interval across the central electrodes (respectively, *p*= 0.001 and *p*= 0.002, corrected for multiple comparisons in space). The topography plots show the contrast of prestimulus HEP amplitude preceding hits and misses between 296–400ms following the R-peak.

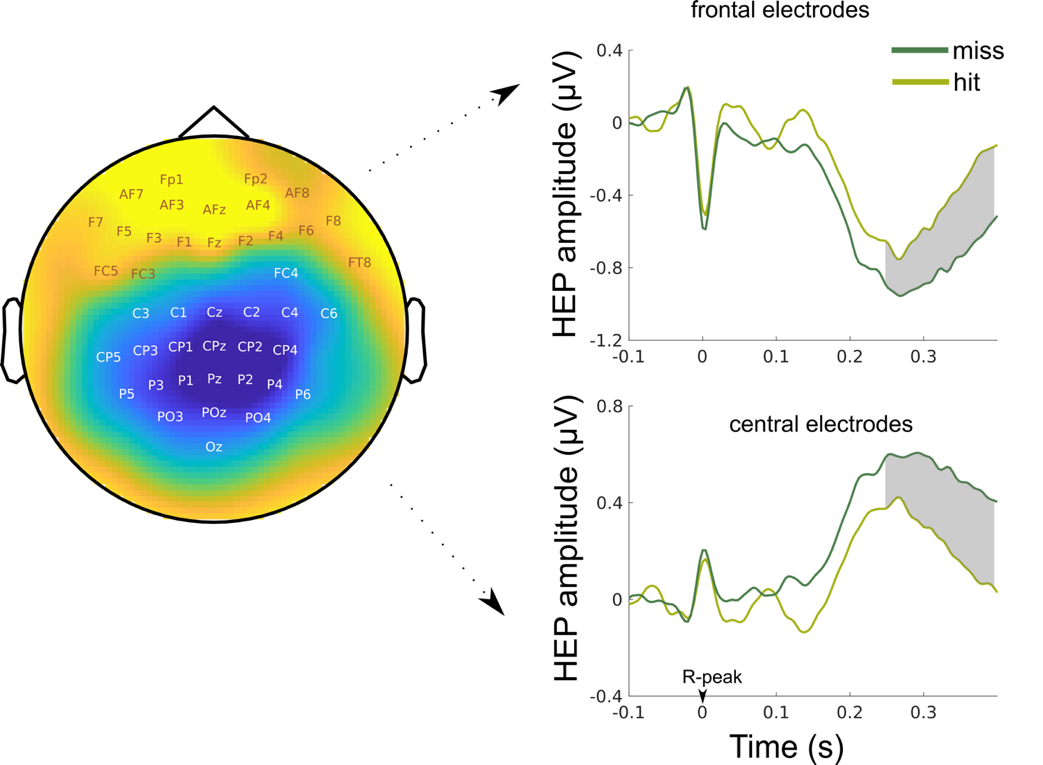

**Supplementary Figure 5.** Exploratory analysis of HEPs amplitudes preceding hits and misses. The topography contrast of prestimulus HEP amplitude preceding hits and misses between 250–400ms following the R-peak. In an exploratory analyses, we also tested overall changes in HEP levels preceding hits and misses across all electrodes for the time window of HEP, 250–400ms after the R-peak. A cluster-based permutation t-test revealed a significant positive cluster over frontal electrodes and a negative cluster over central electrodes between 250–400ms (respectively, Monte-Carlo p=0.001 and p=0.001 corrected for multiple comparisons in space and time). The significant result including negative cluster over central electrodes replicated our hypothesis-driven analyses, where misses were preceded by higher positivity of HEP. The frontal regions in the positive cluster had a negative HEP polarity and the absolute strength of HEP for misses was higher than hits.
